# Human readable compression of GFA paths using grammar-based code

**DOI:** 10.1101/2025.05.22.655470

**Authors:** Peter Heringer, Daniel Doerr

## Abstract

Pangenome graphs offer a compact and comprehensive representation of genomic diversity, improving tasks such as variant calling, genotyping, and other downstream analyses. Although the underlying graph structures scale sublinearly with the number of haplotypes, the widely used GFA file format suffers from rapidly growing file sizes due to the explicit and repetitive encoding of haplotype paths. In this work, we introduce an extension to the GFA format that enables efficient grammar-based compression of haplotype paths while retaining human readability. In addition, grammar-based encoding provides an efficient in-memory data structure that does not require decompression, but conversely improves the runtime of many computational tasks that involve haplotype comparisons.

We present `sqz`, a method that makes use of the proposed format extension to encode haplotype paths using byte pair encoding, a grammar-based compression scheme. We evaluate `sqz` on recent human pangenome graphs from Heumos *et al*. and the Human Pangenome Reference Consortium (HPRC), comparing it to existing compressors `bgzip`, `gbz`, and `sequitur`. `sqz` scales sublinearly with the number of haplotypes in a pangenome graph and consistently achieves higher compression ratios than `sequitur` and up to 5 times better compression than `bgzip` in HPRC graphs and up to 10 times in the graph from Heumos *et al*.. When combined with `bgzip`, `sqz` matches or excels the compression ratio of `gbz` across all our datasets.

These results demonstrate the potential of our proposed extension of the GFA format in reducing haplotype path redundancy and improving storage efficiency for pangenome graphs.

## 1 Introduction

In the past two decades, numerous initiatives have focused on producing reference assemblies for a wide range of species. These reference assemblies play a critical role in tasks such as genotyping, variant calling, and higher-order omics analyses, including epigenomics, metagenomics, and transcriptomics. However, a single reference sequence is insufficient to capture the full genomic diversity within a species and can introduce reference bias into the aforementioned analyses.

In contrast, pangenomes consist of sets of individual haplotype assemblies that collectively represent the genetic variability of a species. Advances in long-read sequencing and assembly technologies have recently enabled the construction of pangenomes from high-quality, reference-grade haplotype assemblies [9, 3] that are particularly suited for identifying structural variation in a species. Although pangenome applications are just emerging, several landmark studies have already demonstrated their advantages in genotyping [16, 2], variant calling [4], and transcriptome analysis [15], outperforming traditional single reference based methods.

In general, as more haplotypes are added to a pangenome, the number of novel genomic variants decreases. A key determinant of the effectiveness of pangenome-based applications is whether the pangenome sufficiently captures the genomic diversity of the species. This is typically assessed by analyzing pangenome growth curves and estimating pangenome openness [19]. These growth curves often plateau, suggesting that a finite set of haplotypes is adequate to represent the genomic variation within a species. Consequently, in the long term, we expect that numerous stable reference pangenomes will be established that will ultimately replace traditional single reference assemblies.

Graph-based data structures have become the predominant approach for representing pangenomes, as they reduce redundancy while preserving properties essential for locating subsequences and visualizing genomic variation. Two widely used graph structures are *sequence graphs* and *de Bruijn graphs*. In sequence graphs, non-overlapping genomic segments are represented as vertices, with edges denoting adjacency relationships between them. These segments are typically obtained from sequence alignments. In contrast, de Bruijn graphs are built by decomposing haplotypes into overlapping sequences of fixed length, called *q*-grams. In this model, *q*-grams form the edges of the graph, and vertices represent the (*q*− 1)-length overlaps between them. In addition to the graph structure itself, pangenome graphs also retain information on the paths of the original haplotype sequences through the graph.

A popular format for storing pangenome graphs is the *Graphical Fragment Assembly* (GFA) format (https://github.com/GFA-spec/GFA-spec), a tabular representation in which each line constitutes a record encoding a specific type of information, such as a node (`S` line), an edge (`L` line), or a haplotype path (`P` or `W` line). The popularity of the format stems largely from its simplicity, human readability, and scripting-friendliness. However, since `P` and `W` lines store path information in an explicit, uncompressed form, file size is typically dominated by these entries that consume excessive amounts of storage, undermining the original advantages of the format. This has led to the ironic situation where, although the size of the pangenome graph itself follows the expected behavior of a saturating growth curve, the corresponding file size increases linearly with the number of haplotypes, as illustrated in Figure 1. For example, the GFA file for a recently released pangenome of human chromosome 19 comprising 1,000 haplotypes occupies 14 GB, whereas the portion encoding the graph structure accounts only for 178 MB. This discrepancy severely hampers the usability of pangenome graphs, as the resulting data volumes become increasingly unmanageable, and it poses a genuine risk to the broader adoption of pangenome-based applications.

**Figure 1:**
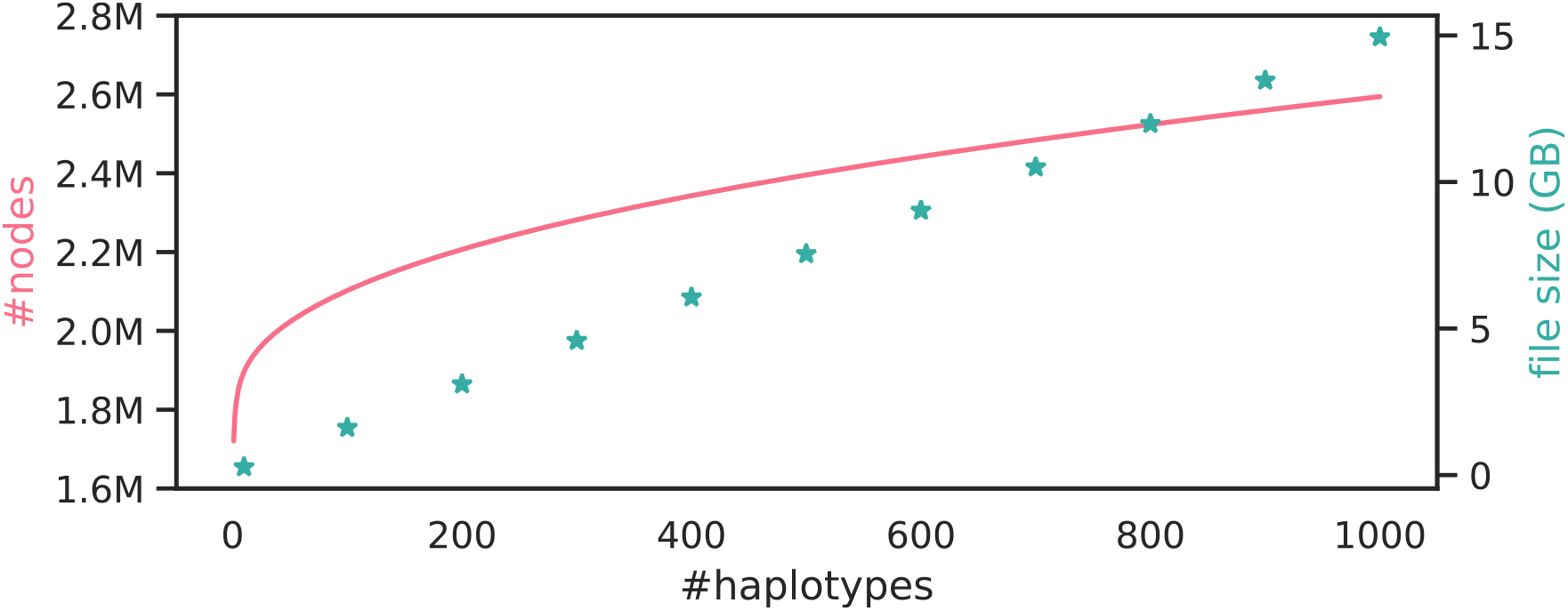
Pangenome growth vs file size. The growth curve computed with Panacus [11] of a pangenome graph of human chromosome 19 comprising 1,000 haplotypes from [5] compared against the file sizes of corresponding subsampled GFA-formatted pangenome graphs. While the growth curve quickly saturates, the file size displays a linear increase.

Currently, to our knowledge only one tool is specifically designed for compressing pangenome graphs: `gbz` [17]. This tool constructs a variant of the *positional Burrows–Wheeler Transform* (PBWT) from the haplotypes encoded in a pangenome graph and also encodes the topology of the sequence graph using succinct data structures. Beyond storage, the `gbz` format provides efficient in-memory representations that support pangenome analysis; for instance, it underpins the `giraffe` read aligner for mapping short reads to pangenome graphs [16]. However, `gbz` is not designed for incremental updates—modifications such as adding nodes or haplotypes require rebuilding the entire structure. The adoption of the `gbz` format within the pangenome graph community has remained limited, largely due to its static and sophisticated data structures, binary file format, and the consequential dependence on external libraries for construction and usage.

In this work, we propose an extension of the more popular GFA file format that allows haplotype paths to be specified in a compressed form, while preserving human readability. Our approach is based on context-free grammars. To this end, we introduce a new line type, the `Q` line, to define *meta-nodes* representing paths in the graph. This enables the representation of repetitive substrings of haplotype paths of length two or more with a single meta-node, resulting in a more compact path encoding. To reference these grammar-defined subpaths within haplotype paths, we introduce an additional line type, the `Z` line. We also present `sqz`, a proof-of-principle implementation, written in Rust, available at https://github.com/codialab/sqz.

There is a long-standing body of research on the concept of *grammar-based code* [6]. In grammar-based coding, nonterminal symbols are introduced to replace subsequences of the original text that occur multiple times. Each nonterminal is associated with a production rule that maps it to a sequence of symbols–potentially including other nonterminals–that ultimately spells the original subsequence. A prominent grammar-based coding scheme is *byte pair encoding* (BPE), in which each production rule corresponds to a digram, i.e., a pair of adjacent symbols, that appears more than once in the text. To construct the grammar, BPE iteratively identifies the most frequent digram, replaces all of its occurrences with a new nonterminal symbol, and records the corresponding production rule. This process continues until no digram appears more than once in the transformed text. BPE and its variants, such as SentencePiece [7], have become standard machine learning, particularly in large language model (LLM) applications, where compact and consistent tokenization is crucial for managing vocabulary size and handling rare or unseen words efficiently [13, 12].

## 2 Methods

### 2.1 Preliminaries

A *sequence graph G* = (*V, E*) is an undirected graph that represents DNA molecules drawn from an alphabet Σ and links between them. A DNA molecule is composed of two antiparallel strands of oligonucleotides, with one being designated *forward* and the other *reverse complementary* strand. DNA molecules are represented by the vertices of the graph *G*, with each vertex *v* in *V* having a *left* and a *right* side. A visit of vertex *v* must respect the direction, that is, if *v* is entered on its left side, it must be exited to its right side, and vice versa. If a node is traversed left-to-right, then it spells the forward strand of the corresponding DNA molecule, while a right-to-left traversal produces the corresponding reverse complementary strand. Any two (not necessarily distinct) vertices can be connected through an undirected edge. To indicate on which ends the two vertices connect, we mark vertices traversed in right-to-left orientation with an overline, e.g., 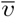 with the default (non-overlined) reading direction being left-to-right. For instance, if the left side of vertex *u ∈ V* is connected to the left side of vertex *v ∈V*, then 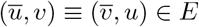. We will also use the short form of *uv* to refer to edge (*u, v*).

A *haplotype path*, or simply called *haplotype*, is a walk through *G*. Note that a haplotype can also be a cycle or cover only a single vertex, although in the following, we will consider without loss of generality that all haplotypes start and end with distinct vertices. A *pangenome* is a tuple (*G, H*) constituting a sequence graph *G* = (*V, E*) and a set of haplotypes *H* that covers *G*, that is, each edge in *E* is traversed at least once by any haplotype of *H*. In incomplete assemblies, haplotypes may be split in smaller contigs, leading to a haplotype being associated with multiple paths. For presentation purposes, we adopt a simplified notation assuming each haplotype is represented by a single path, however, this does not diminish the generality of our approach.

### 2.2 Extension of the GFA format

Reflecting the nature of the existing GFA format, we formulated several design requirements for a compressed representation of haplotypes that extends this format:

- *Human readable*. The format should remain accessible and interpretable by humans, rather than resembling machine code.
- *Simple*. The compressed information should be intuitively understandable when browsing a GFA file, consistent with the style of existing record types.
- *Updatable*. It should be straightforward to add new nodes and edges to the graph, as well as new compressed haplotype path encodings without requiring decompression or modification of previously stored compressed paths.
- *Versatile*. The format should accommodate various compression strategies rather than being tied to a single algorithm.

We propose a grammar based coding scheme for haplotype compression by introducing two new record types, whose fields are detailed in Table 1. The `Q` record defines a *meta-node* representing a reusable walk within the pangenome graph. These meta-nodes can be defined recursively and may include references to other `Q` records. For example, the `W` record associated with sample `S1` in Figure 2d, which defines the haplotype path (1, 4, 6, 8, 9) in the graph shown in Figure 2a, can be encoded as (1, *q*_2_, 9) using the `Q` records *q*_1_ = (4, 6) and *q*_2_ = (*q*_1_, 8). To store such compressed paths, we introduce the `Z` record, which maintains the same structure as the `W` record but supports references to meta-nodes. The resulting compact encoding of all `W` records shown in Figure 2d is illustrated in Figure 2e.

**Table 1:**
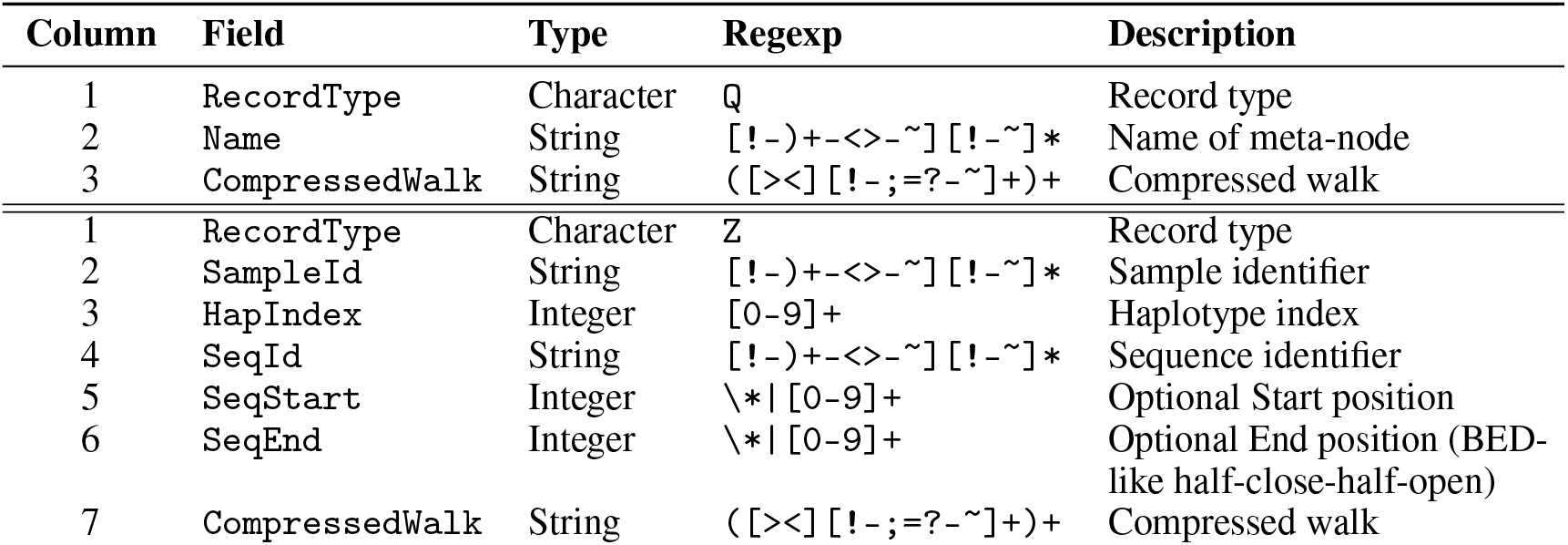
Definitions for Q and Z lines that enable grammar based coding in GFA format. The record definitions are analogous to those described in https://github.com/GFA-spec/GFA-spec/blob/master/GFA1.md.

**Table 2:**
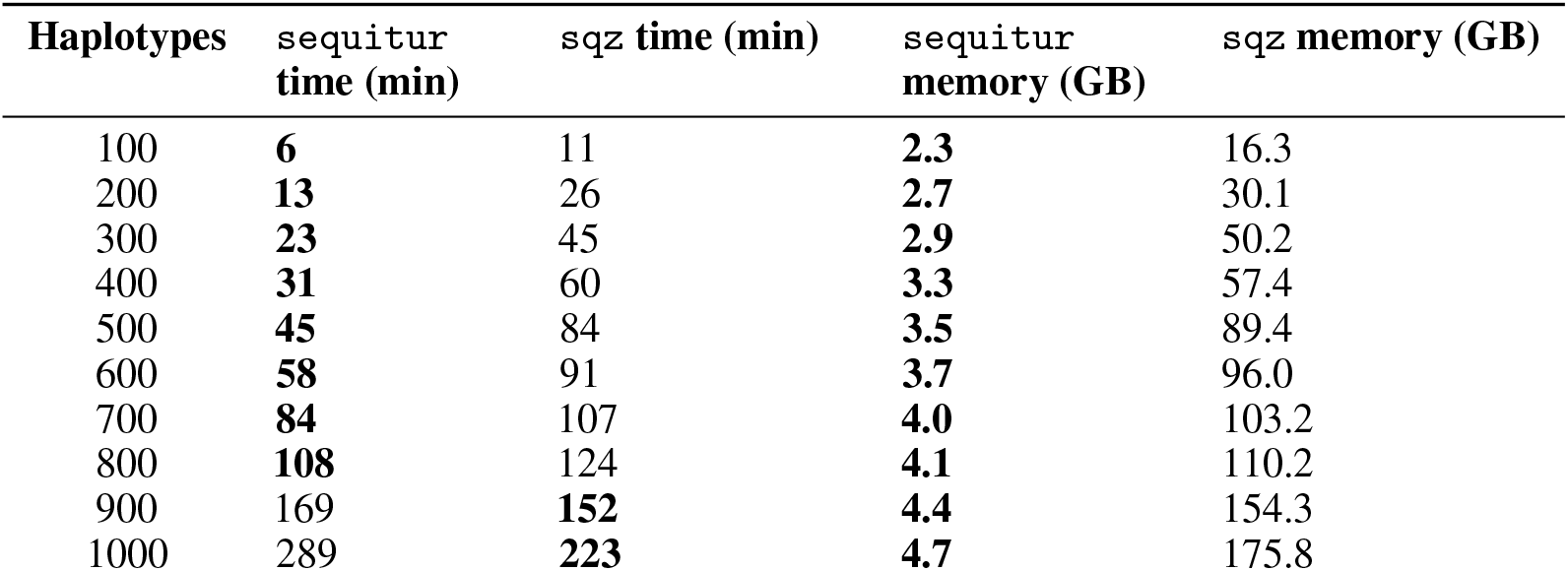
Runtimes and peak memory usage of sequitur and sqz on the chromosome 19-1000 graph with different numbers of haplotypes.

| Haplotypes | sequitur<br>time (min) | sqz time (min) | sequitur<br>memory (GB) | sqz memory (GB) |
| --- | --- | --- | --- | --- |
| 100 | **6** | 11 | **2.3** | 16.3 |
| 200 | **13** | 26 | **2.7** | 30.1 |
| 300 | **23** | 45 | **2.9** | 50.2 |
| 400 | **31** | 60 | **3.3** | 57.4 |
| 500 | **45** | 84 | **3.5** | 89.4 |
| 600 | **58** | 91 | **3.7** | 96.0 |
| 700 | **84** | 107 | **4.0** | 103.2 |
| 800 | **108** | 124 | **4.1** | 110.2 |
| 900 | 169 | **152** | **4.4** | 154.3 |
| 1000 | 289 | **223** | **4.7** | 175.8 |

**Figure 2:**
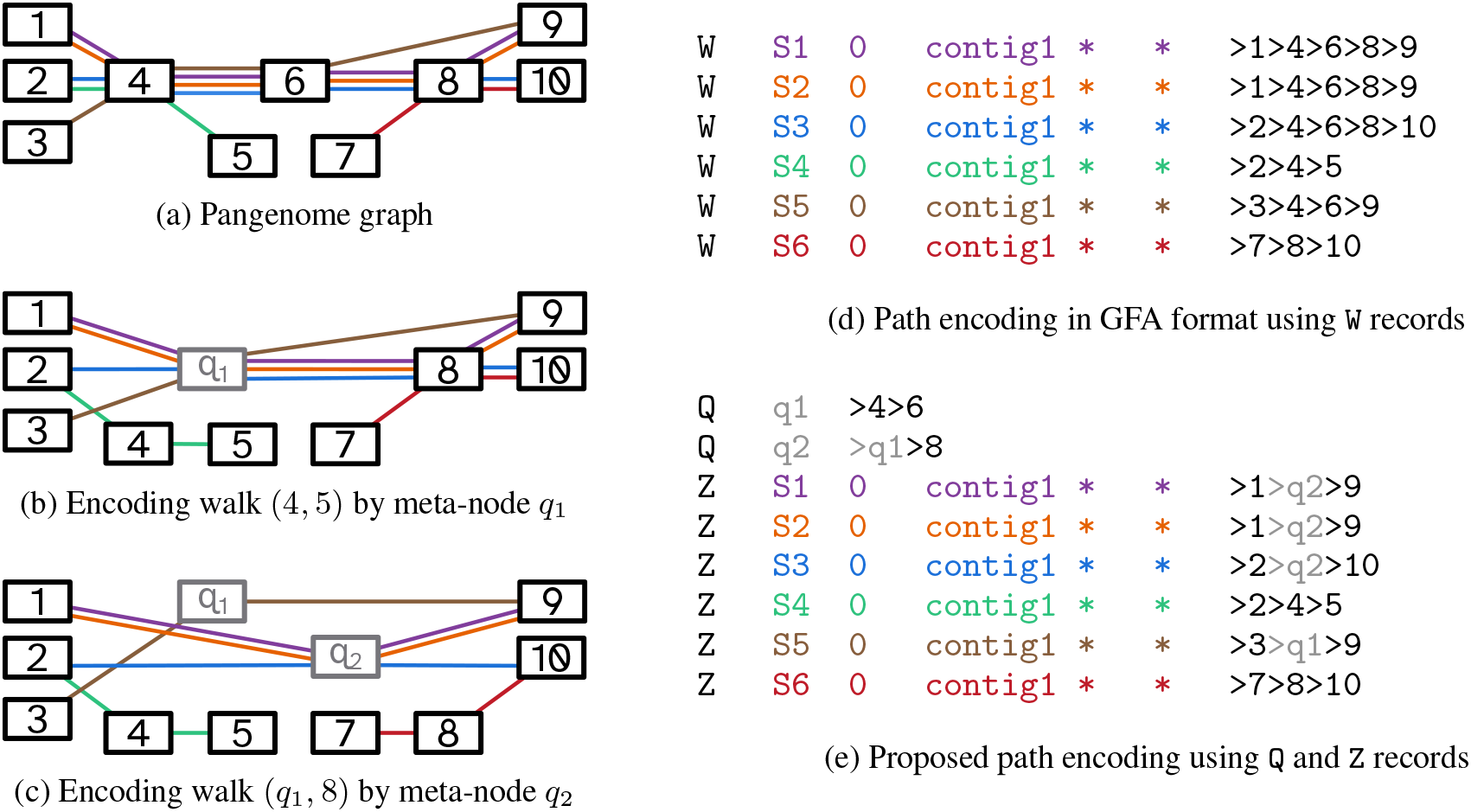
Haplotype compression with Q and Z records. **(a)** A pangenome graph containing 6 haplotypes. **(b)** The repeated walk (4, 6) is replaced by meta-node *q*_1_. **(c)** The repeated walk (*q*_1_, 8) is replaced by meta-node *q*_2_. **(d)** Original `W` records for each haplotype. **(e)** Compressed encoding using `Q` and `Z` records. Note that repeated walks (1, *q*_2_) and (*q*_2_, 9) could be replaced by additional meta-nodes.

### 2.3 Grammar constructing using byte pair encoding

Our method is composed of three steps that will be explained in the following sections:

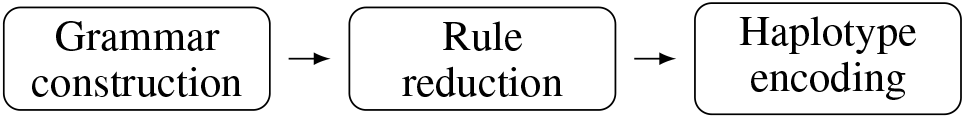

For grammar construction, we adopt a variant of *byte pair encoding* (BPE). BPE iteratively replaces frequently occurring *digrams*, i.e, pairs of adjacent symbols, with new nonterminal symbols and corresponding production rules that reconstruct the original sequence. At each iteration, the most frequent digram is selected for replacement, with ties broken arbitrarily.

Unlike natural language text, haplotype paths in pangenome graphs are *bidirected*, meaning they can be traversed in both forward and reverse-complementary direction. In our adaptation, digrams correspond to pairs of adjacent oriented vertices in the pangenome graph, and repeats are detected irrespective of orientation. Because haplotypes may revisit the same vertex multiple times, maintaining the correct ordering of digrams is crucial—especially since the algorithm processes digrams by frequency rather than by their original order in the haplotype paths. To preserve sequential information, we annotate each digram with a pair of increasing integers, which we refer to as an *address*:

#### Definition 1.

*An address is a pair of non-negative integers α* | *β associated with a digram, satisfying α* ≤ *β. Given a haplotype H represented as a sequence of successive digram occurrences, H* = *h*_0_ · · · *h*_*m*_, *the addresses of consecutive digrams are overlapping: that is, for all* 0 ≤*i < m, if h*_*i*_ = (*wu, α* | *β*) *and h*_*i*+1_ = (*uv, γ* |*δ*), *then β* = *γ. Additionally, each digram occurrence in H is unique within the haplotype*.

One way to construct addresses is a left-to-right traversal of each haplotype such that the first number of the address of a digram matches the second number of the preceding address:

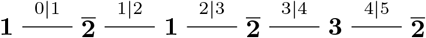

As shown, the digram 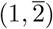 occurs twice. By assigning addresses, we retain the relative position of each digram, enabling accurate identification of its neighbors—even when digrams are processed out of sequence.

Our algorithm for grammar construction, presented in Algorithm 1, begins by initializing table *D* with all digrams observed across the haplotypes. Each edge *uv* in the pangenome graph is associated with an entry *D*[*uv*], which stores a list of tuples (*i, a*_1_ |*a*_2_) indicating that digram *uv* appears in haplotype *H*_*i*_ with address *a*_1_ |*a*_2_ (see line 1). The algorithm proceeds by iteratively replacing the most frequent digram with a new meta-node, continuing until no digram occurs more than once (see line 3). Replacing a digram *uv* with a meta-node *q* introduces two new digrams, *wq* and *qw*′, where *wu* and *vw*′ are the digrams immediately preceding and succeeding *uv* at each of its occurrences across the haplotypes. Accordingly, occurrences of *wu* and *vw*′ that are embedded in a walk of the form *wuvw*′ are removed from the corresponding lists *D*[*wu*] and *D*[*vw*′], and the updated digrams *wq* and *qw*′ are added to new entries *D*[*wq*] and *D*[*qw*′], respectively. These updates are handled by the for-loops in lines 6 and 14, which generate the entries *D*[*wq*] and *D*[*qw*′] by traversing the occurrences of the currently handled digram and finding their neighbors. To ensure that the sequential structure of digrams is preserved, the addresses of the new digrams must be reconciled. This is done in line 16: the first number of each address in *D*[*qw*′] is updated to match the first number of the corresponding entry in *D*_*q*_, which, by construction, equals the second number of the address in *D*[*wq*] (see Figure 3a).

**Figure 3:**
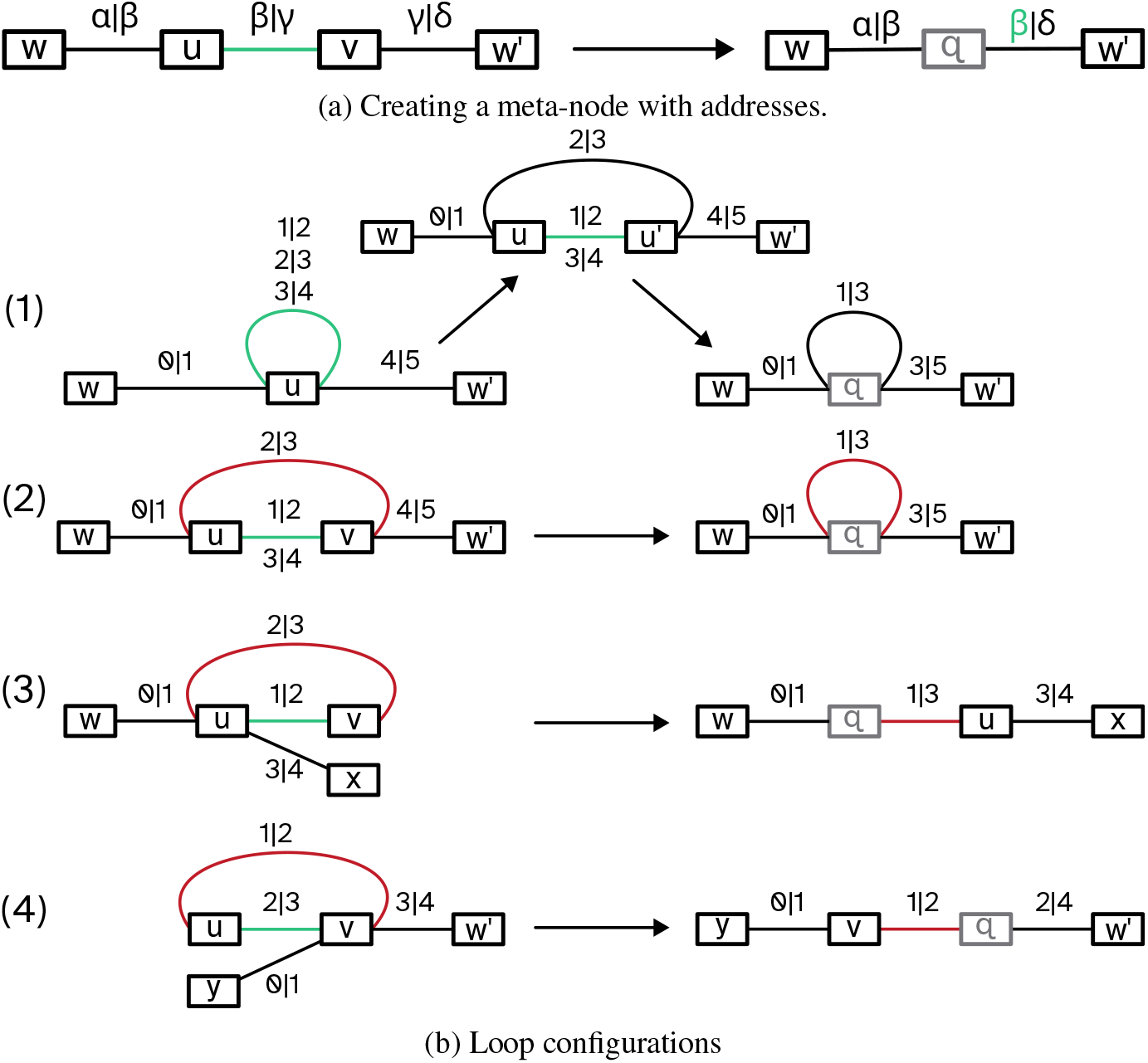
**(a)** During meta-node creation neighboring addresses need to be changed to keep the address guarantees that neighboring digrams share a number. **(b)** A number of special cases. (1) is a self loop digram that needs to be split into two edges before a meta-node can be created. In case (2) a meta-node creation along the green edge results in the creation of a new self loop based on the red edge. Case (3) and (4) appear similar to (2) yet do not result in self loop edges.

Our algorithm has to accommodate four special loop configurations that are treated within the conditional block at line 10 and illustrated in Figure 4a. The first occurs when *u* = *v* and *u* forms a loop on its own, while the remaining three configurations occur when distinct (meta-)nodes *u* and *v* together form a loop:

**Figure 4:**
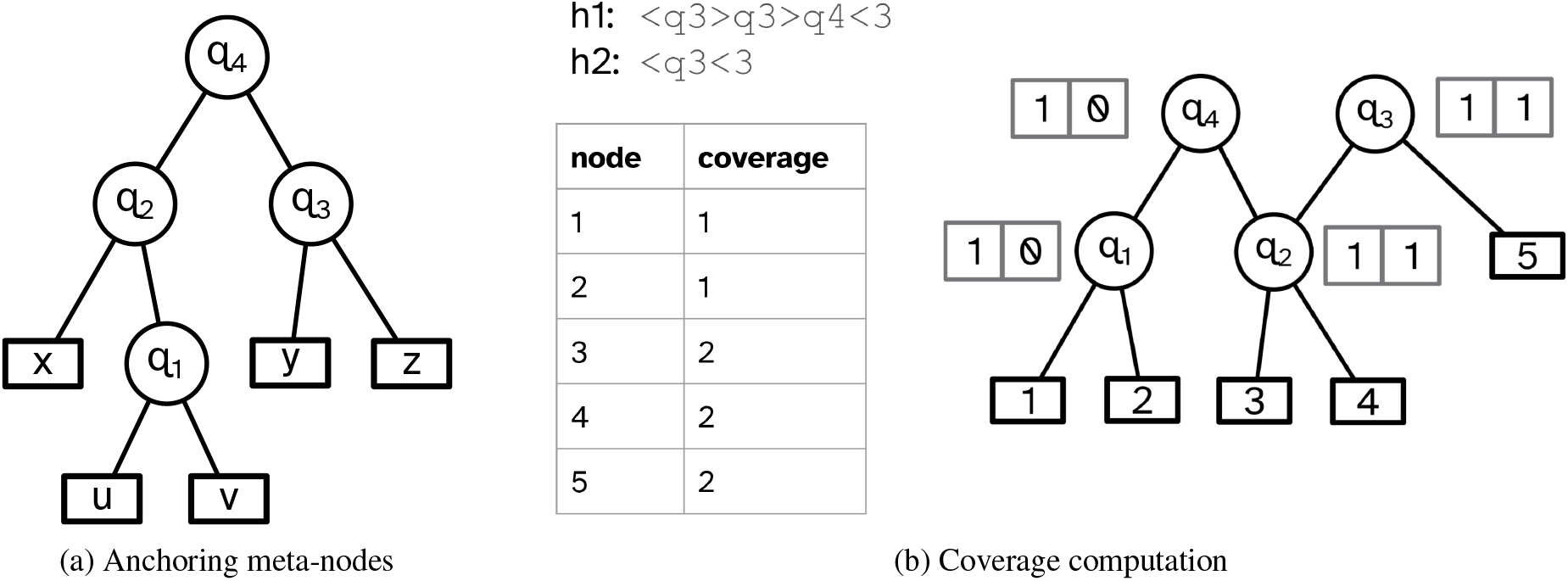
**(a)** For the encoding of haplotypes the leftmost and rightmost anchoring meta-nodes have to be found. Additionally, the offset between the leftmost (rightmost) anchoring meta-nodes and the true leftmost (rightmost) child is needed. For *q*_4_, *q*_1_ is the leftmost anchoring meta-node with an offset of 1 (due to node *x*) and *q*_3_ is the rightmost anchoring meta-node. **(b)** Coverages of nodes in haplotypes can be calculated by counting nodes in compressed paths and creating presence/absence lists. These lists are then pushed down the directed acyclic graph of the grammar to get the final lists. The coverage for a vertex is the number of presence entries in its list.

1. *u* = *v (self-loop), with u appearing in a walk of the form w****uu****uw*′. At least two consecutive addresses are associated with digram *uu* (green edge) of which every second must be transferred onto the new meta-node. This is handled by extracting every second address and placing them into a temporary list *D*[*uu*′] in line 11 that is then processed in the conditional block along with the next cases.
2. *uv appears in a walk of the form w****uv****uvw*′. Any address of the reverse arc *vu* (red edge) that simultaneously matches two addresses of *uv*, one by its first number and the other by its second, must be transferred into *D*[*qq*], forming a new self-loop at the meta-node.
3. *uv appears in a walk of the form w****uv****ux*. If an address of arc *vu* matches only the second number of an address of *uv*, it is moved to *D*[*qu*] and does not contribute to a new self-loop.
4. *uv appears in a walk of the form yv****uv****w*′. If an address of arc *vu* matches only the second number of an address of *uv*, it is moved to *D*[*vq*] and does not contribute to a new self-loop.

*Runtime*. For a given pangenome graph *G* = (*V, E*) with haplotypes *H*_1_, …, *H*_*k*_, the total number of digram occurrences across all haplotypes is given by 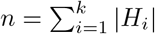. Accordingly, the size of the digram table *D* after initial construction in line 1 is also 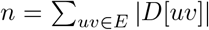. We implement *D* as a hash table. Since we know the maximum number of entries of the table is *n*, we can pre-allocate a large enough table, which provides constant-time access, insertion, and removal of entries. The total number of digram occurrences *n* remains constant throughout the algorithm: digram occurrences are neither created nor deleted, but are merely redistributed among entries in *D* (as in the loops at lines 6 and 14). In each iteration, the algorithm processes a set of *x* = |*D*[*uv*] | digram occurrences and then removes the entry *D*[*uv*] (line 18). Hence, the runtime can be captured by the recurrence *T* (*n*) = *T* (*n* − *x*) + *f* (*x*), where *f* (*x*) accounts for retrieving and updating all data related to *D*[*uv*]. With appropriate auxiliary data structures, we ensure that *f* (*x*) ∈ *O*(*x*).

#### Algorithm 1 Grammar construction

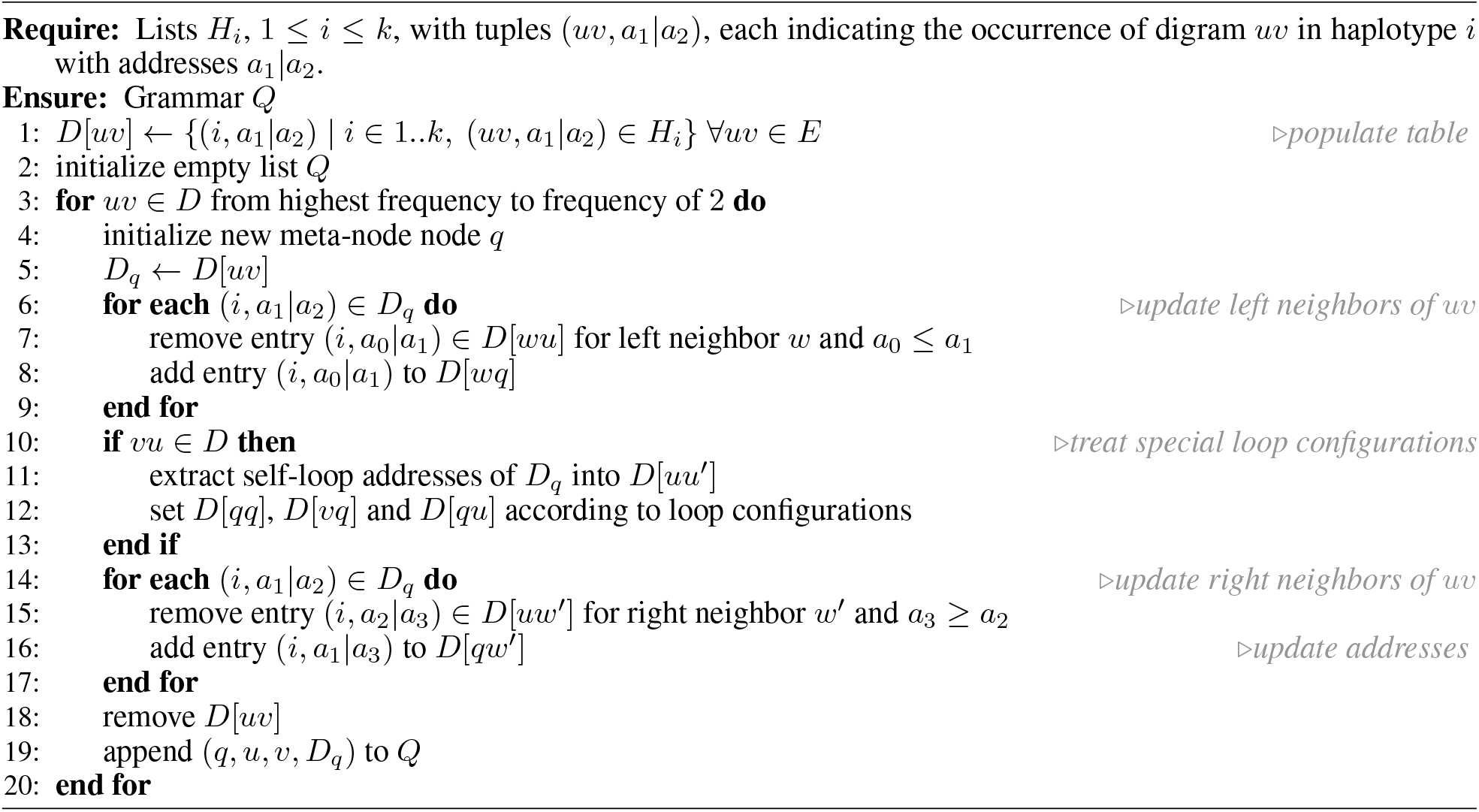

We first introduce a frequency-based structure, Freq, to process digrams in linear time by descending frequency. Let *m* = max_*uv* ∈*E*_ |*D*[*uv*] | be the maximum initial digram frequency. We implement Freq as a list of *m* entries, where index *i* stores a hash set of digrams with frequency *i* (for 2 ≤ *i* ≤ *m*). Initialization of Freq requires a single pass over *D* and runs in *O*(*n*) time. During the main loop (line 3), digrams are processed in decreasing order of frequency using Freq. As new digrams are created, their frequencies are computed and added to the appropriate buckets in Freq. Specifically, new digrams of the form *wq* and *qw*′, created in the for-loops at lines 6 and 14, are inserted into Freq after computing their respective frequencies |*D*[*wq*] | and |*D*[*qw*′]|. Importantly, the frequency of any newly created digram cannot exceed the frequency of the currently processed digram *uv*.

Two additional hash tables, NeighborLeft and NeighborRight, track adjacency relationships between (meta-)nodes in haplotypes. For each occurrence (*i, a*_1_ |*a*_2_) of a digram *uv*, we store an entry NeighborRight[(*u, i, a*_1_)] = (*v, a*_2_) which points to the succeeding (meta-)node *v* and its associated number *a*_2_. Analogously, NeighborLeft[(*v, i, a*_2_)] = (*u, a*_1_) points to the preceding (meta-)node *u* and its associated number *a*_1_. These structures allow constant-time retrieval of address components, as in lines 7 and 15. The removal of (*i, a*_0_|*a*_1_) from *D*[*wu*] and insertion into *D*[*wq*] then takes constant time. Simultaneously, we update

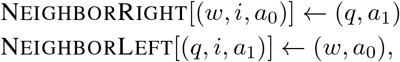

and remove the entry NeighborLeft[(*u, i, a*_1_)]. All operations are in constant time. Analogous updates occur in line 15. The space complexity is *O*(*n*), as all data structures, *D*, Freq, NeighborLeft, and NeighborRight, are bounded by the number of digram occurrences. This completes the runtime analysis. Since *f* (*x*) ∈ *O*(*x*), applying the substitution method to the recurrence *T* (*n*) yields an overall linear runtime.

### 2.4 Space improvements

Haplotypes corresponding to chromosome-level assemblies can contain hundreds of millions of nodes in pangenome graphs. As a result, address numbers must be stored as 64-bit integers, which leads to high memory usage. We now present an encoding scheme for addresses that scales with the number of repetitions in a sequence rather than with sequence length. This allows both numbers of an address to be stored together within a 64-bit representation, effectively halving the memory required.

We achieve this by splitting each haplotype into segments. Each segment is defined by a node appearing twice, except for the last segment, where the right boundary is determined by the end of the sequence. In doing so, segments overlap by one position, called the *pivot* that constitutes the second appearance of the repeated node in the left segment. Note that the first appearance of the repeated node can be at any preceding position of the segment:

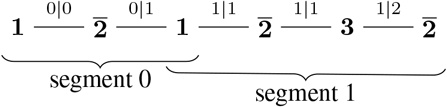

Each segment is assigned a dedicated value that determines both numbers of the addresses of all digram occurrences within that segment except for the digram immediately preceding the pivot that connects to the next segment. This special digram occurence must preserve the overlapping property of addresses as defined in Definition 1, ensuring that neighboring addresses overlap. In the example above, the segment with the dedicated value 0 is linked to its succeeding segment with value 1 by a transition digram carrying the address 0|1, positioned just before their shared pivot.

#### Proposition 2.

*A digram appearing multiple times on the same haplotype can always be differentiated by its addresses*.

*Proof*. By construction, in each segment only one node can appear twice and therefore no digram occurrence is repeated.

#### Proposition 3.

*Addresses of consecutive digram occurrences overlap*.

##### Algorithm 2 Haplotype encoding

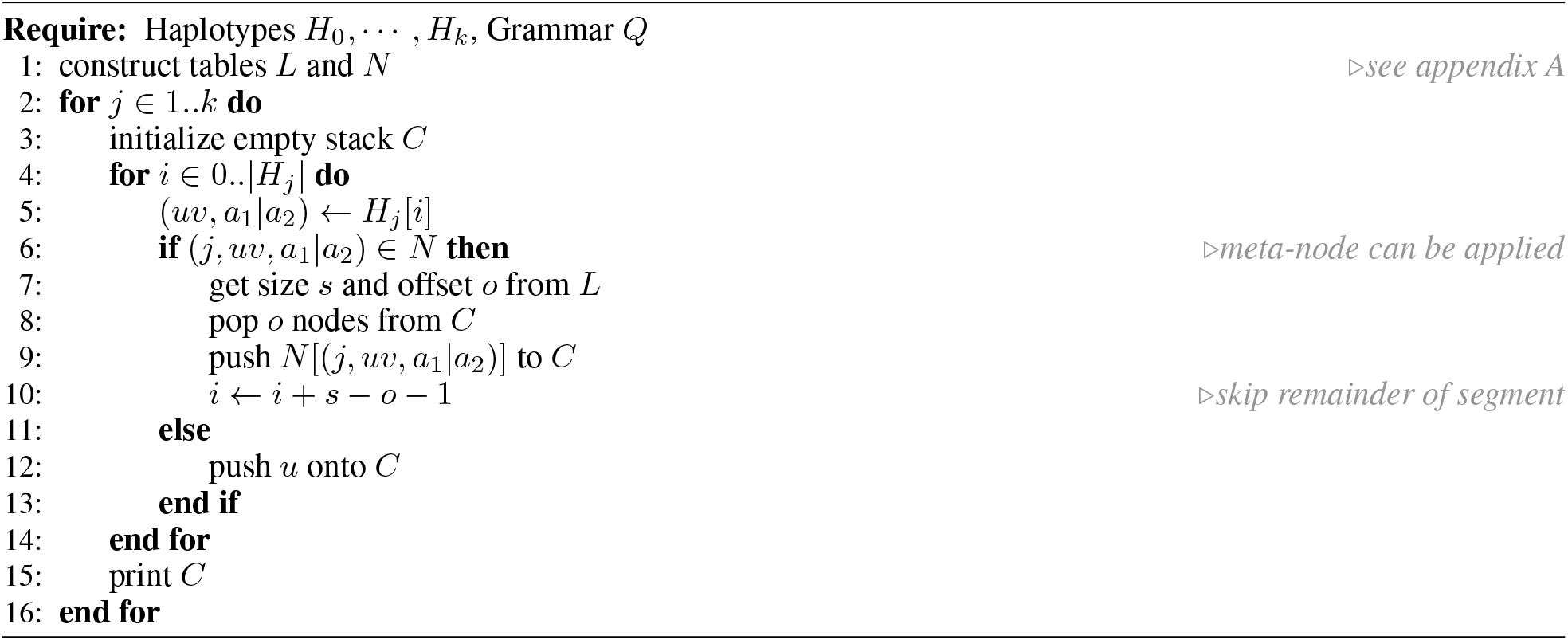

*Proof*. All addresses of a segment composed of the dedicated value overlap. The address of the segment’s last digram, which occurs left of the pivot, is composed of the dedicated values of the current and the next segment, again producing an overlap. □

### 2.5 Reducing grammar rules

Before encoding, we remove meta-nodes that appear only once, either in the set of production rules (`Q` lines) or in the haplotype encoding (`Z` lines). A meta-node is removed by replacing it with the outcome of its corresponding production rule, after which this rule is discarded from the grammar. To identify such candidates, we first count for each meta-node the number of production rules in which it is used. To identify how often a meta-node is used in the haplotype encoding, we count the number of digram occurrences associated with each meta-node. A decrease in this count between a meta-node and its parents indicates that the meta-node also appears in at least one haplotype encoding. We denote such nodes in the following as *top-level meta-nodes*.

### 2.6 Haplotype encoding

In the following, we present a linear-time algorithm for the encoding of haplotypes from the constructed grammar. Our approach relies on the identification of *anchoring meta-node*s associated with top-level meta-nodes. Anchoring meta-nodes are nodes whose two children are leaves, that is, they constitute pairs of vertices in the pangenome graph. Each top-level meta node, corresponding to an entire segment of one or more digram occurrences, has a left-most and right-most anchoring meta-node. However, not every right-most and left-most meta-node of a top-level meta-node is an anchoring meta-node, as illustrated in the example of Figure 4a.

Our algorithm, shown in Algorithm 2, produces the encoding of a haplotype without repeated steps of inserting and replacing intermediary meta-nodes. Each haplotype is processed separately in the main loop (line 2). The algorithm then iterates over each position *i* associated with a digram occurrence (*uv, a*_1_ | *a*_2_) in the current haplotype *H*_*j*_ (line 4). If (*j, uv, a*_1_ |*a*_2_) is associated with an *anchoring meta-node*, the entire segment is replaced by its top-level meta-node (line 6). Otherwise, node *u* is pushed onto the stack. This is necessary as some nodes of the haplotype–those not being contained in the grammar–will not be replaced by any meta-node.

We utilize two auxiliary data structures that allow us to determine a top-level meta-node from a single digram occurrence.

1. Table *L* stores for each meta-node *q* its size (in nodes), its leftmost and rightmost anchoring meta-node, and corresponding *offsets*. The offsets correspond to the number of leaf vertices within *q* that are left of its leftmost anchoring meta-node and respectively right of its rightmost anchoring meta-node. This table is used in the construction of the following table.
2. Table *N* maps digram occurrences of anchoring meta-nodes to their top-level meta-node.

Table *L* is constructed in a bottom-up traversal of the directed acyclic graph constituting the grammar. In each step, one of the following two cases applies to the currently processed meta-node *q*:

1. Meta-node *q* is already an anchoring meta-node, and therefore, its leftmost and rightmost anchoring meta-node is *q* itself. Accordingly, *q* is stored in *L* with size 2, corresponding to the number of its children, and offsets 0.
2. Otherwise, we determine *q*’s size, its left-most and rightmost anchoring meta-node, and their offsets by looking up *q*’s children in *L*.

We then construct table *N* by iterating over all top-level meta-nodes, and creating for each such meta-node *q*′ entries in *N* that map digram occurrences associated with its leftmost and rightmost anchoring meta-node to *q*′ itself.

#### Runtime

In the following, we study the runtime of our algorithm under a grammar where each production rule maps to exactly two (meta-)nodes, as it is the case prior to rule reduction. This will simplify the analysis, which operates on the directed acyclic graph structure underlying the grammar, but does not change its outcome: If we remove a meta-node *q*, then all the edges to *q*’s children are moved to *q*’s parent. Additionally, the edge between *q* and its parent is removed. The total numbers of edges and nodes are reduced by one, respectively. Therefore, we can analyze without loss of generality the grammar before reduction, as it becomes only smaller through the reduction process.

Since we know the sizes of *L* and *N* in advance, we can pre-allocate the exact amount of memory, allowing us *O*(1) insertions. The construction of *L* has to iterate over all the meta-nodes of the grammar. The lookup of the children of each meta-node is performed in constant time by querying *L*. Thus, the runtime of *L*’s construction is limited by *O*(|*Q*|).

To construct *N*, again all the rules are iterated, however this time also all of its digram occurrences are traversed. If the meta-nodes are processed in top-down order, we can simply create a hash set of all the digram occurrences whose top-level meta-nodes were already visited. Then the check whether a node is a top-level node can be done in *O*(1). Since all of the remaining instructions are also constant-time look-ups, the runtime is bounded by the number of total digram occurrences, which is *O*(*n*).

Haplotype encoding is performed by iterating over all digram occurrences. Inside the loop, each iteration adds at most one element to the stack. This means we can pop only as much elements as we have iterations. Additionally, the final stack for each haplotype has at most the length of the inner loop, meaning the print instruction has the same complexity as the inner loop. Therefore the whole runtime is depending only on the number of digram occurrences, limiting the worst-case runtime to *O*(*n*).

### 2.7 Coverage computation with compressed haplotypes

In this section, we provide evidence that the compressed grammar can serve as an efficient in-memory data structure and speed up haplotype analysis while simultaneously reducing memory usage. To this end, we address the task of computing a node coverage table. In this table each vertex of the pangenome graph is recorded with the number of haplotypes in which it appears at least once.

For this task, the grammar-encoded haplotype paths do not need to be decompressed. In fact, the calculation can be even more efficient by exploiting the directed acyclic graph that underlies the grammar. We iterate over the encoded haplotypes and produce presence/absence lists for all the (meta-)nodes, similar to how this process is done on uncompressed haplotypes. However, since the compressed haplotypes are shorter than uncompressed counterparts this process is faster. In the next step, the presence/absence lists are pushed down the directed acyclic graph using a top-down approach, as seen in Figure 4b. Lists from parental meta-nodes are combined using disjunction. Thereafter, lists of the leaf nodes are collected and summed to produce the reported coverage table.

## 3. Results

We implemented our method, `sqz`, in Rust and released the source code under the MIT license at https://github.com/codialab/sqz.

We evaluated `sqz` on three human pangenome graphs:

- *Chr19*, comprising 1,000 haplotypes [5],
- *HPRC v2*.*0 MC GRCh38 full* and
- *HPRC v2*.*0 MC GRCh38 clipped*, both comprising 464 haplotypes [9].

The full version of the HPRC v2.0 MC GRCh38 graph includes complete assemblies, but alignment quality is poor in certain regions. Particularly centromeres contain essentially large chunks of unaligned sequence. In contrast, the clipped version removes these segments, limiting the pangenome graph to regions with reliable alignments.

We benchmarked our method against three compression tools: `bgzip` [1], `gbz` [18], and `sequitur` [10]. `bgzip` produces `gzip`-compatible compressed binary files and is widely used for compressing pangenome graphs in the GFA format, primarily because its output can be decompressed with `gzip`, which is pre-installed on most Unix systems. Its key advantage over `gzip` is support for indexing, allowing selective decompression of file segments. `gbz`, in contrast, is specifically designed for compressing pangenome graphs and integrates with tools in the GBWT ecosystem, as discussed in Section 1. `sequitur` is a classical grammar-based compression algorithm in natural language processing (NLP).

Among existing BPE based algorithms, Re-Pair [8] is most similar to our approach. Both algorithms iteratively replace the most frequent digram with a production rule or meta-node, proceeding in descending order of frequency. However, Re-Pair operates on linear text rather than graph structures, and it does not account for inverted digrams, unlike `sqz` and `gbz`. Although several Re-Pair implementations are available, many operate solely on text or binary files and are not designed to handle characters spanning multiple bytes. Therefore, we were unable to successfully apply them to haplotype paths, which are composed of node identifiers that cannot be represented by a single byte.

In contrast, we were able to evaluate `sequitur` [10], another BPE algorithm. `sequitur` constructs and applies production rules as soon as it reads a repeated digram. As a result, `sequitur`’s memory usage scales with the size of the compressed output rather than the input. However, its compression performance depends on the input order, since it processes digrams in the order they appear rather than by frequency. `sequitur` allows the user to set the size of its core data structure, a hash table. This affects both memory usage and runtime: larger tables reduce collisions and improve performance at the cost of increased memory. For all evaluations, we set the hash table size to 2,000 MB, which was the maximum value supported by the program.

As an initial experiment, we used the *Chr19* pangenome to evaluate how compression scales with the number of haplotypes in a pangenome graph. To this end, we randomly removed haplotypes until only a specified number remained, and removed all S- and L-lines corresponding to nodes and edges that are not covered by the remaining subset. We then applied `sqz`, `bgzip`, `sequitur`, and `gbz` to the resulting trimmed graphs. Additionally, we tested a combined approach using `sqz` followed by `bgzip` compression, as `bgzip` is commonly used in transferring GFA files.

The resulting file sizes and relative space savings are shown in Figure 5. Space saving is defined as 1− (compressed*/*uncompressed). `bgzip` exhibits constant compression behavior: file size increases linearly with the number of haplotypes, and space saving remains constant at about a rate of 0.74. For fewer than 100 haplotypes, `bgzip` achieves better compression than the purely grammar-based approaches `sqz` and `sequitur`. For all other methods, the compression rate improves as more haplotypes are added, benefiting from increased redundancy. Up to 100 haplotypes, the difference between `sqz` and `sequitur` remains small (e.g., 0.19 GB vs. 0.24 GB), but it gets larger with increasing number of haplotypes. Among all methods, `gbz` and the `sqz+bgzip` combination deliver the best compression results, with `sqz+bgzip` slightly outperforming `gbz` for fewer haplotypes.

**Figure 5:**
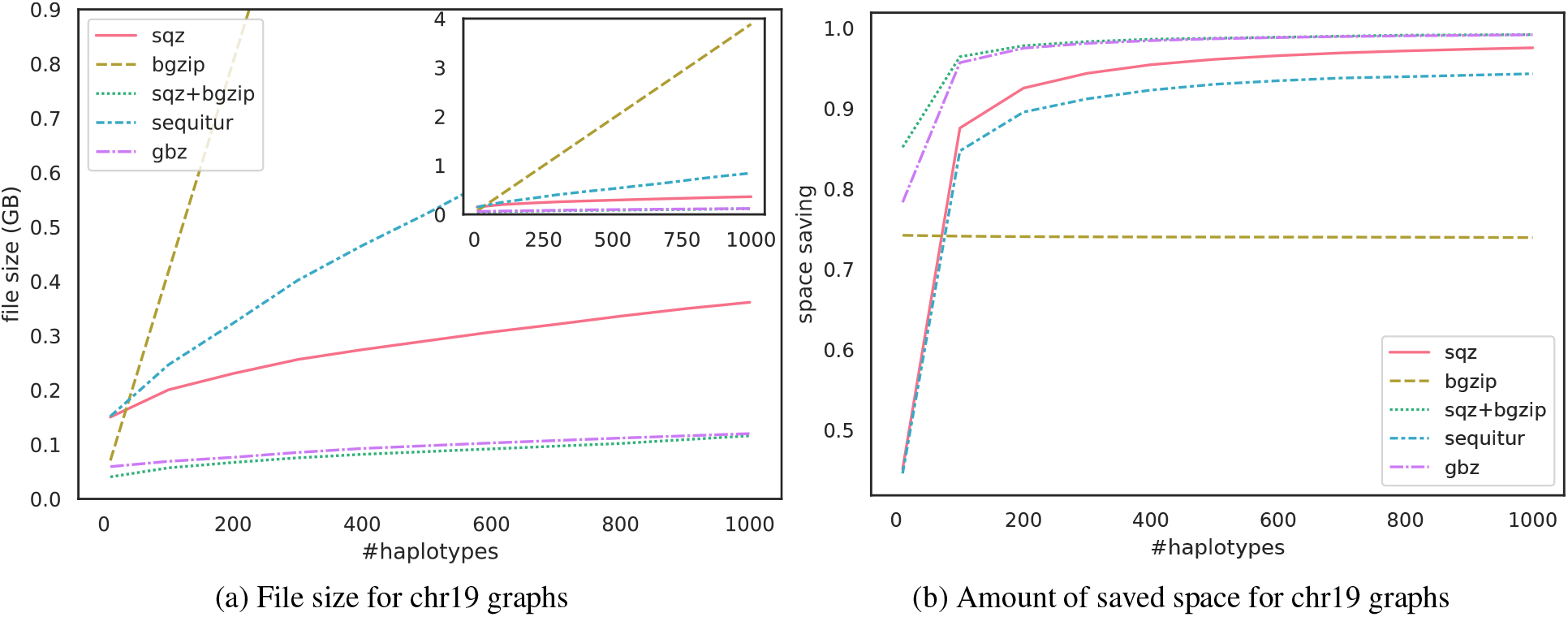
File sizes and amount of saved space for chr19 subgraphs with only the specified number of haplotypes included.

A second benchmark was conducted on the clipped and full HPRC v2.0 MC GRCh38 graphs, that have been split up into their individual chromosome graphs. This setup enables an evaluation of how compression performance is affected by the similarity among haplotypes, due to the full graphs containing less similar haplotype paths than their clipped counterparts.

Due to the large input sizes (up to 36 GB), we were unable to run `sequitur` for this benchmark, as it required excessive runtimes. We evaluated compression performance in terms of both file size and compression ratio, defined as uncompressed*/*compressed. For the clipped graphs, `sqz+bgzip` and `gbz` perform similarly again, achieving compression ratios between 60 and 80, depending on the specific graph. `sqz` alone achieves a compression ratio of around 20, while `bgzip` performs worst overall. Except for `bgzip`, compression results on the full graphs vary considerably, as the amount of heterochromatic sequence differs across chromosomes. For chromosomes 1 through 11, `bgzip` typically outperforms `sqz`, but from chromosome 12 onward, `sqz` yields smaller files, with the exception of chromosomes 1 and 9. `gbz` consistently produces smaller files than `bgzip`, while the combined `sqz+bgzip` approach generally results in the smallest file sizes, again with the exception of chromosome 9.

To assess how effectively haplotypes are compressed using `sqz`, we examined the number of nodes per haplotype for each chromosome in the clipped graphs, illustrated in Figure 7. `sqz` achieves a reduction in haplotype length by more than two orders of magnitude. This analysis also reveals a non-negligible variance across haplotypes, with some chromosomes, e.g. chromosome 15, containing haplotypes that are much less compressed than others.

**Figure 6:**
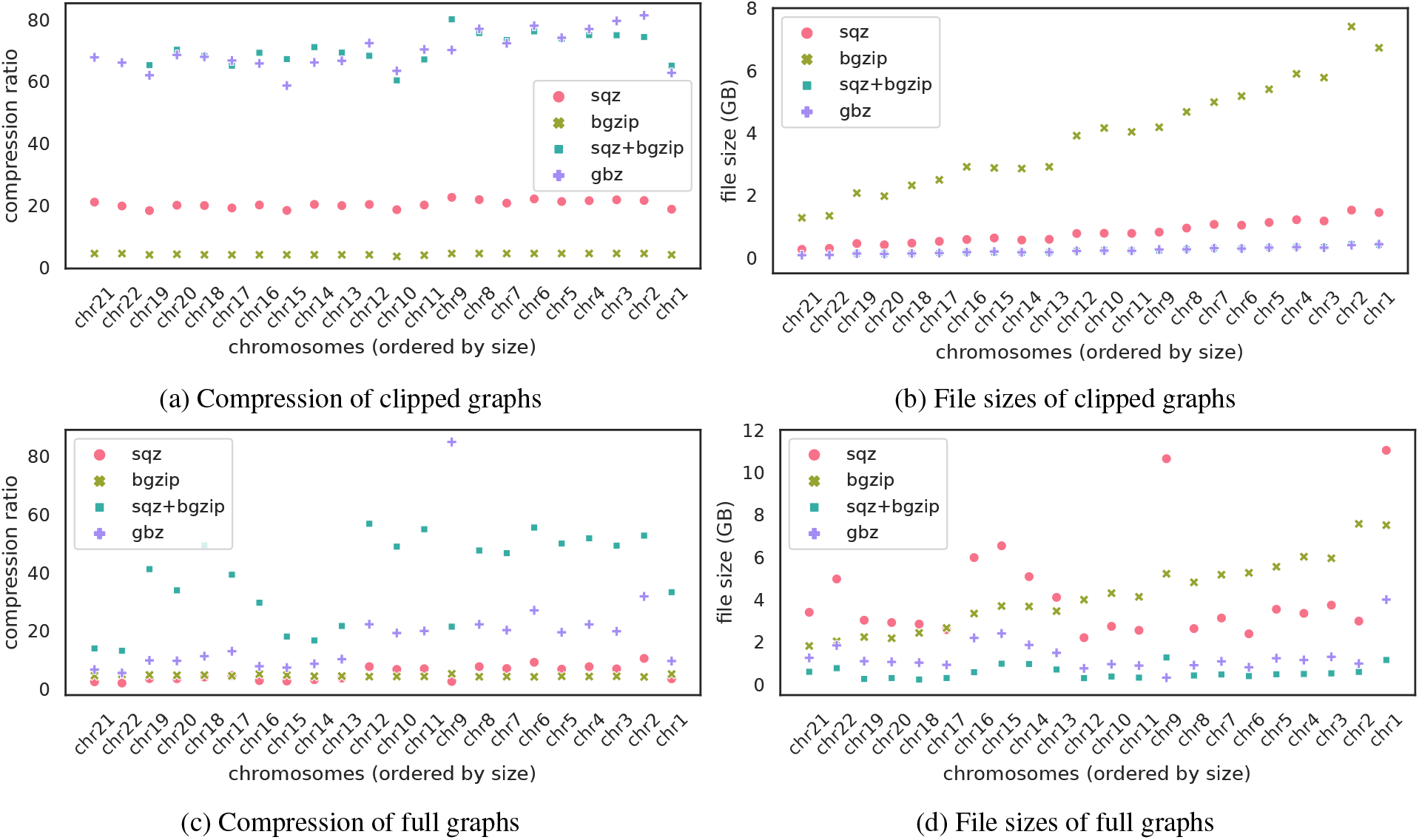
Compression of the HPRC v2.0 MC graphs

**Figure 7:**
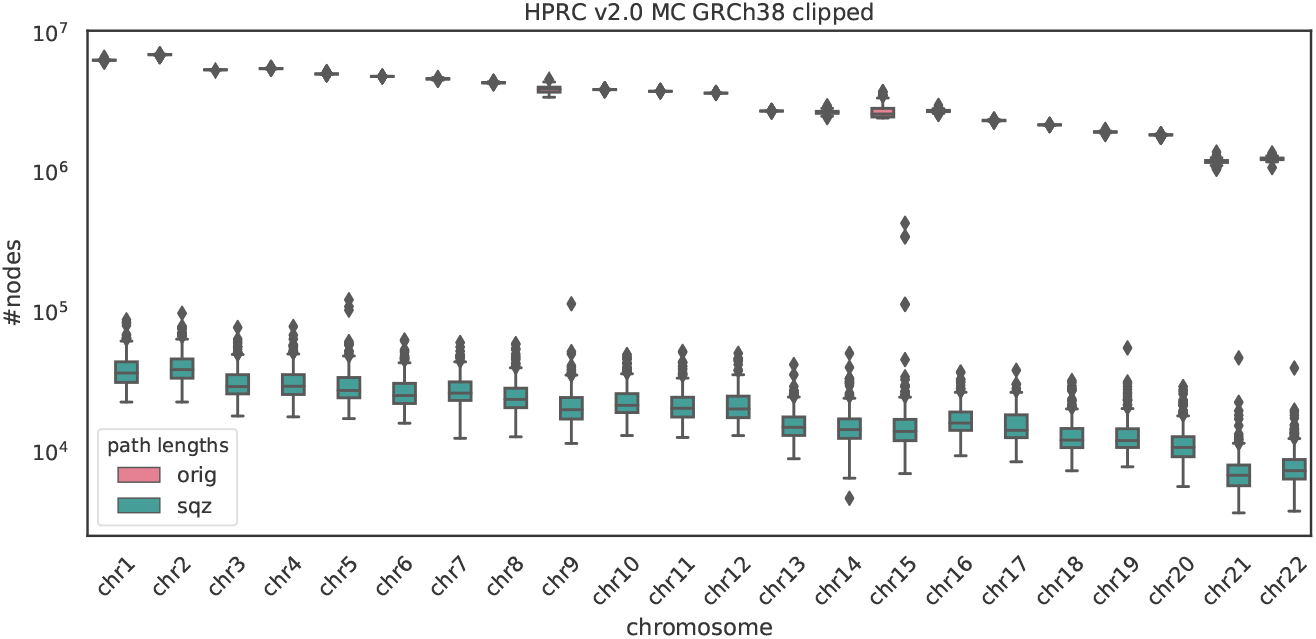
Number of node entries per haplotype per chromosome

Another key consideration when working with large files is the runtime and memory consumption of the compression algorithm. We compared the performance of `sequitur` and `sqz` on the Chr19 pangenome using the set of trimmed graphs containing a varying numbers of haplotypes that we constructed for the first experiment. Only these two tools were included in this comparison, as `bgzip` and `gbz` follow fundamentally different compression strategies.

In terms of runtime, `sequitur` outperforms `sqz` on graphs with up to 800 haplotypes. However, for the two largest graphs with 900 and 1,000 haplotypes, `sqz` is faster and, for the latter graph, requires more than an hour less runtime. On memory usage, `sequitur` outperforms `sqz` by a wide margin. Even for the largest graph, `sequitur` stays below 5 GB of memory usage, while `sqz` requires up to 176 GB.

To evaluate the performance of computing coverage tables using the in-memory representation of the grammar, we implemented two Python scripts that make use of libraries `networkx` and `numpy`. One does the calculation on an uncompressed GFA file of the Chr19 pangenome. The second one follows the approach described in the previous section and takes a GFA file compressed by `sqz`. The script for the uncompressed GFA file takes 7:49 min, while the one on the `sqz`-compressed graph only takes 2:39 min. The memory usage is reduced over ten-fold when using the script with the compressed graph, requiring only 14.8 GB instead of 159.6 GB. We also ran the two approaches over all of the HPRC v2.0 graphs (clipped and full), the results are shown in Figure 8. Similar to the Chr19 graph, working with the grammar-based strategy is faster and uses less memory for all chromosomes. Particularly, the difference in terms of runtime is large, with the original method needing 2:08 hrs, while the compressed files only need 12 min for the clipped chromosome 1.

**Figure 8:**
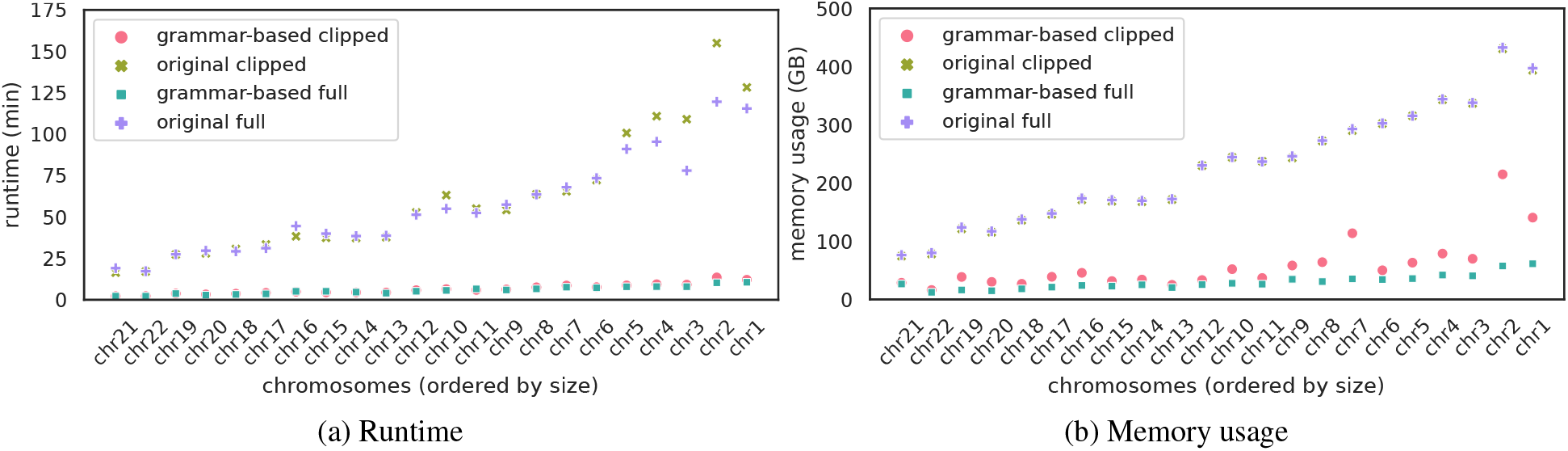
Coverage calculation on clipped and full HPRC v2.0 MC GRCh38 graphs.

## 4 Discussion and Outlook

By introducing Q- and Z-records, we propose an extension to the GFA format that supports human-readable, grammar-based compression. Grammar-based compression is simple and intuitive, yet is as effective as any finite-state compression scheme [6]. Our encoding is also incrementally updatable: nodes and edges can be added to the graph without interfering with the grammar captured in Q-records. Likewise, new haplotypes can be encoded using the existing grammar, although this may lead to less efficient compression. Grammar-based encoding is highly versatile and has given rise to a broad body of research, resulting in a variety of compression schemes [6]. We also envision further variants, such as grammars based on *q*-grams for *q >* 2, or grammars derived from *maximal exact matches* (MEMs) which are widely used in pangenomics [14]. Moreover, Q-records can carry biological meaning by annotating shared sequences associated with specific alleles. This enables not only compression, but also allows to encode haplotypes as combinations of alleles, embodying a natural, biologically motivated representation.

Most importantly, we argue that grammar-encoded haplotypes need not be decompressed for analysis. Instead, they enable faster and more memory-efficient haplotype analysis, as discussed in the previous section.

Our BPE-based implementation, `sqz`, serves as a proof of principle for the proposed GFA extension. `sqz` often achieves higher compression ratios than `bgzip` and `sequitur`, and performs on par with `gbz` when combined with `bgzip`.

However, `sqz` offers several advantages over `gbz`: it provides a simple, human-readable, and incrementally updatable encoding. Additionally, the resulting grammar can serve as a highly efficient in-memory data structure, without the need for external libraries. Its simplicity allows it to be implemented using built-in data structures available in widely used scripting languages such as Python.

## A Auxiliary data structures for haplotype encoding

### Algorithm 3 Create table *L*

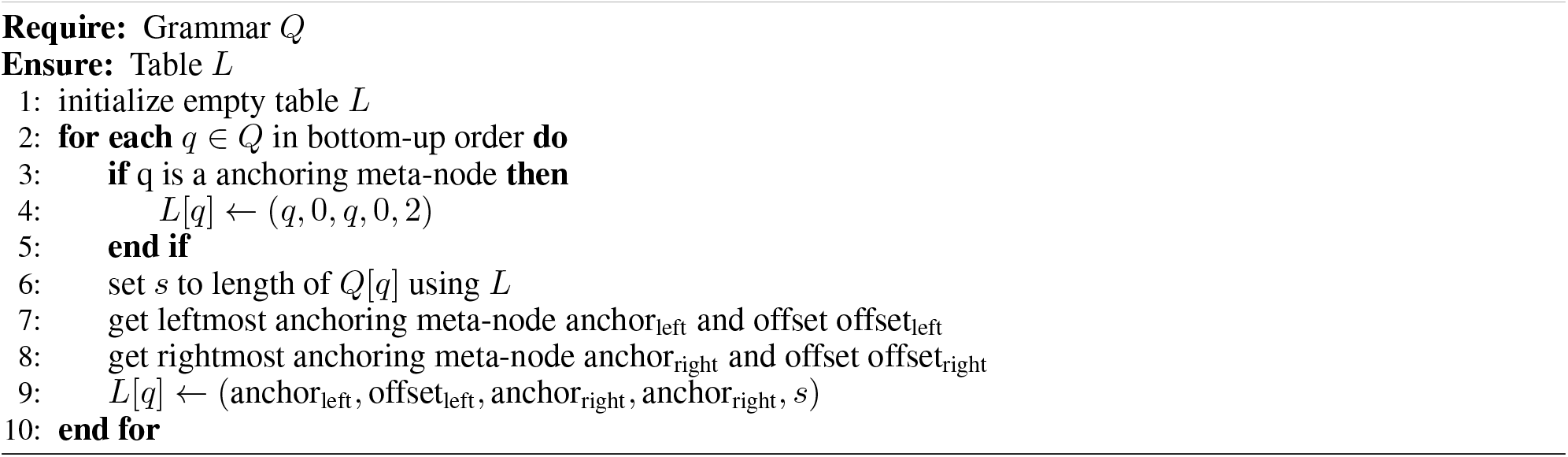

### Algorithm 4 Create table *N*

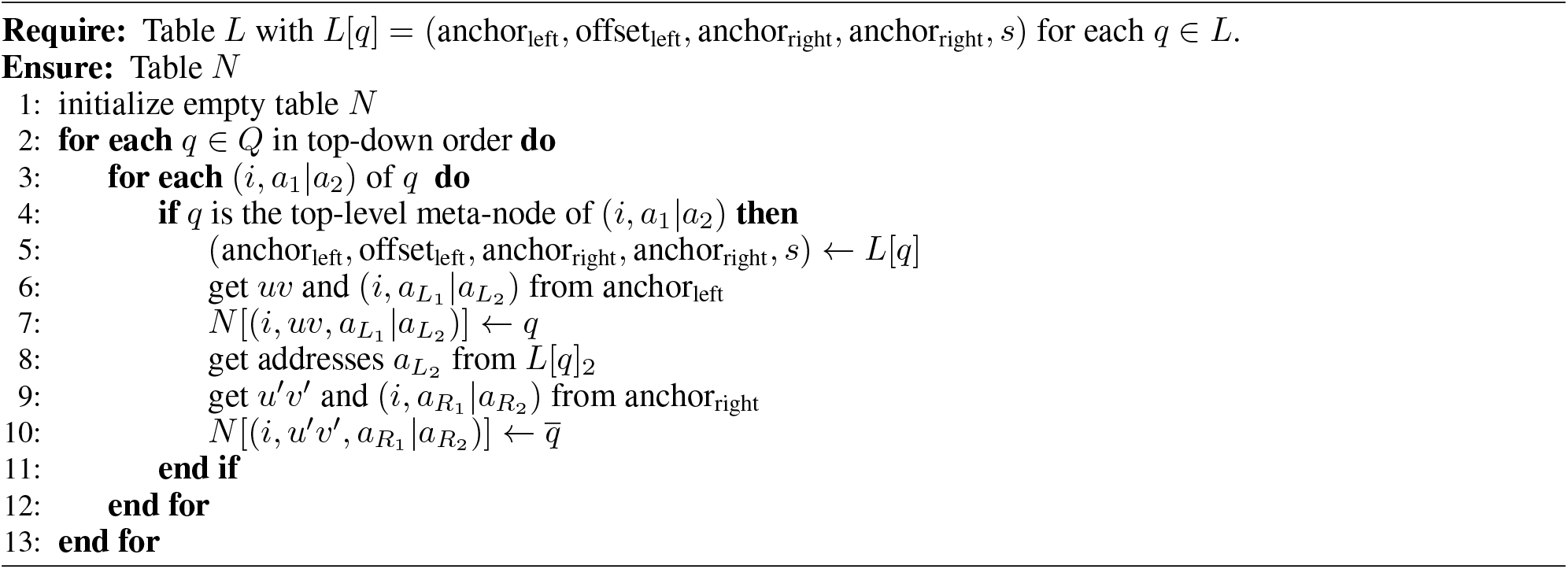

